# Biomechanics of static roll posture control by body flexion in adult zebrafish

**DOI:** 10.1101/2025.05.26.655680

**Authors:** Ryota Nagaoka, Taisei Katayama, Shin-ichi Higashijima, Masashi Tanimoto

**Affiliations:** Division of Behavioral Neurobiology, National Institute for Basic Biology, Okazaki, Aichi 444-8787, Japan; Neuronal Networks Research Group, Exploratory Research Center on Life and Living Systems, Okazaki, Aichi 444-8787, Japan; The Graduate University for Advanced Studies, SOKENDAI, Okazaki, Aichi 444-8787, Japan; Division of Biological Science, Graduate School of Science, Nagoya University, Nagoya, Aichi 464-8602, Japan

## Abstract

Posture control is crucial for animals. Static posture control in fish remains poorly explored. Recent studies have shown that larval zebrafish perform body flexion during slight roll tilts: the body flexion displaces a gas-filled swim bladder, generating counter-rotation torque for postural recovery through misalignment of gravity and buoyancy forces. Swim bladder deflation impairs this postural recovery, suggesting its critical role. This static posture control strategy may be utilized by many fish species. However, adult fish differ from larvae in morphology and behavior, raising questions about the generality of this mechanism. Our behavioral analysis showed that adult zebrafish also flexed their body during roll tilts, with flexion persisting until recovery to an upright posture. Similarly to larvae, swim bladder deflation impaired postural recovery. These results demonstrate that adult fish employ static roll posture control through body flexion and suggest the generality of this mechanism in fish species.

## Introduction

For many animals, posture control is crucial for survival, regardless of whether they live on the land or in the water. Terrestrial animals use both dynamic and static postural control mechanisms.^1–3^ When posture is significantly disrupted, animals dynamically move their limb and axial muscles to recover their desired posture. Even during seemingly motionless periods, static postural control through anti-gravity muscle tone continuously maintains posture.

How about postural control in fish? Previous studies have been predominantly centered on dynamic movements involving swimming and fin dynamics.^4–8^ In contrast, our understanding of static postural control mechanisms remains limited. This is particularly intriguing because most fish species are inherently unstable^7^ - anesthetized or dead fish typically roll to a belly-up position due to their mass distribution.^6,9–11^ Nevertheless, many fish species maintain an upright posture even while appearing motionless.^9,10^ These common observations suggest the existence of static postural control mechanisms through neuromuscular activity. However, until recently, the underlying mechanisms of this static postural control remained poorly understood.

Recent studies on millimeter-sized larval zebrafish have provided valuable insights into this longstanding question. When slightly roll tilted or artificial vestibular stimulation mimicking a head roll tilt was applied, larval zebrafish perform body flexion and recover an upright posture.^12–16^ This body flexion behavior is driven by neural circuits, from a vestibular nucleus via reticulospinal neurons projecting to the spinal cord, and the posterior hypaxial muscles (PHM) near the swim bladder.^12,17^ This posture control through body flexion relies on the presence of the swim bladder, a gas-filled chamber in the body: postural recovery is impaired in swim bladder-deflated larvae.^12^ Biomechanics underlying this postural recovery has been proposed.^12^ The body flexion during roll tilt displaces the swim bladder to one side while placing the head and tail to the other side. Since the density of the swim bladder is orders of magnitude lower than the rest of the body, this asymmetric positioning of the swim bladder results in the misalignment of gravity and buoyancy forces, which generates a corrective moment of force for postural recovery. This fine and static posture control strategy may be utilized by many fish species possessing a swim bladder or low-density tissue. However, adult fish differ from larvae in musculoskeletal structure, body size, and behavioral dynamics,^18–24^ raising questions about the generality of this mechanism.

In the present study, behavioral analyses reveal that adult zebrafish utilize body flexion to correct roll-tilted posture similarly to larvae. When sustained roll tilt stimulus is applied, the body flexion persists throughout the stimulus. Crucially, swim bladder deflation disturbs postural recovery, suggesting swim bladder’s critical role in maintaining static posture. These results suggest that zebrafish maintain an upright posture via the fine and static roll posture control mechanism through body flexion throughout their life.

## Results

### Adult zebrafish utilize body flexion to correct roll-tilted posture

To investigate how adult zebrafish maintain an upright posture, we developed a fish rotation apparatus (Figure 1A). A fish was placed in a cylindrical chamber in a dark room and subjected to postural disturbance in the roll axis. Viscous water containing methylcellulose was used to efficiently deliver roll stimulus to the fish and to slow down its behavior for easier observation. Fish’s posture was simultaneously recorded by two cameras positioned at the front and dorsal sides of the chamber. Pigment-less *casper*^25^ fish were used as a standard fish for easier deflation of the swim bladder in later experiments (see below).

**Figure 1.**
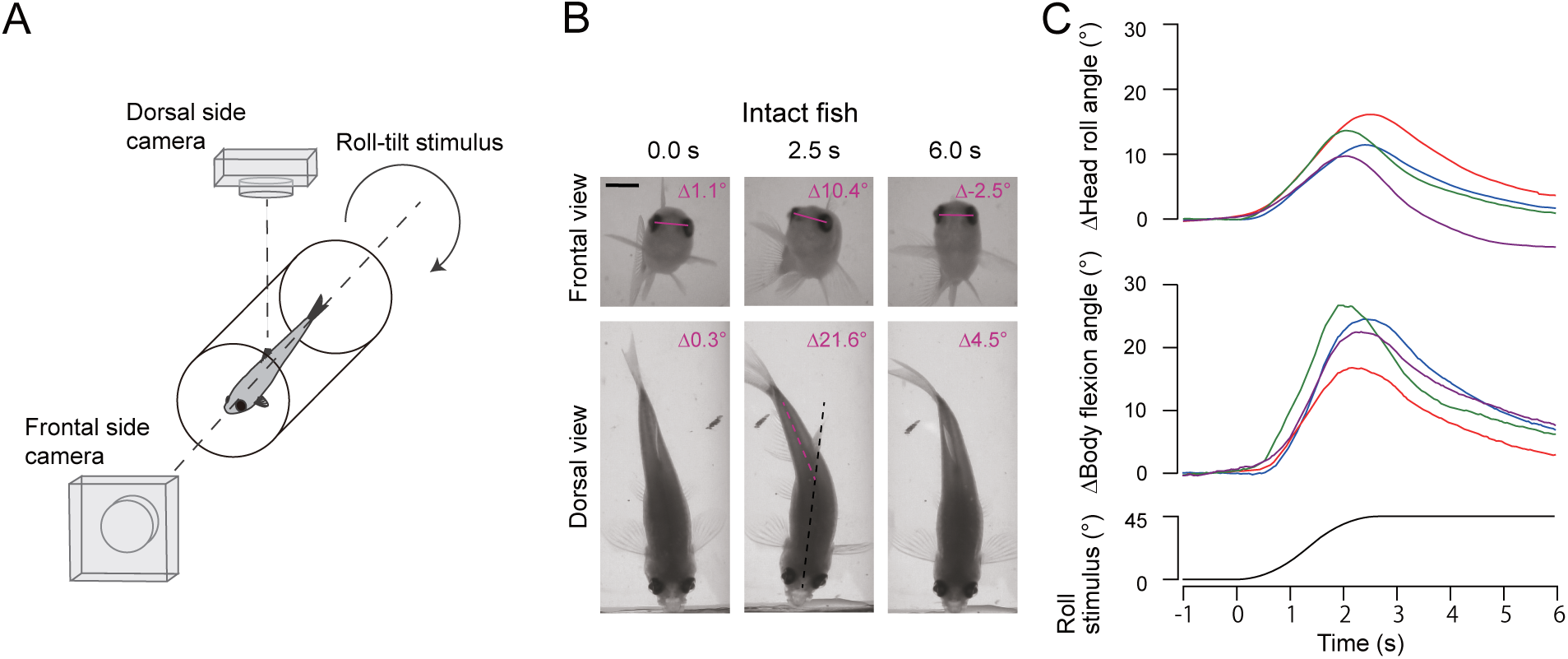
Static roll posture control through body flexion in adult zebrafish. (A) Schematic illustration of unrestrained behavioral experiments. Behavior of adult zebrafish during roll tilt stimuli was recorded using dorsal and frontal view cameras in a dark room. (B, C) Behavioral experiments on intact adult *casper* fish. (B) Snapshots of a fish during a roll tilt stimulus towards its left-down side (top row: frontal view; bottom row: dorsal view). Magenta lines connecting the center of eyes show head roll angle. Black and magenta dashed lines denote the midlines of the rostral and caudal body, respectively. Changes in the head roll angle and body flexion angle relative to the pre-stimulus baselines are indicated at the upper right. Fish tank water contains methylcellulose. Before stimulus onset, the fish maintains an upright posture (0.0 s). As the fish is tilted in the roll direction, it flexes its body towards the ear-up side (2.5 s). The fish recovers its upright posture as it straightens its body (6.0 s). Scale bars indicate 3 mm. (C) Time courses of changes in head roll angle and body flexion angle in response to roll tilt stimuli. Average traces from four individual fish are shown in different colors (7 to 9 trials per fish). See also Figures S1.

Rotation of the chamber induced roll tilt in the fish. As previously reported in larvae,^12,13,15^ adult fish occasionally performed dynamic postural control behaviors accompanied by swim-like vigorous body movement and fin beating. Since the present study focused on static postural control behaviors, we collected trials in which fish recovered posture without these dynamic movements. When a roll-tilt stimulus was applied towards fish’s left-down side, fish was slightly tilted to the left-down direction. Simultaneously, the fish flexed its body to the right (ear-up) side (Figure 1B middle column, Video S1). As the tilt stimulus progressed, head roll angle and body flexion angle increased (Figure 1C). After the roll tilt stimulus reached its final angle, fish gradually recovered the upright posture, while the flexed body relaxed to its original state (Figure 1B right column, Figure 1C, Video S1). Similar results were obtained in wild-type fish (Figure S1). These results suggest that upon postural disturbance in the roll axis, adult zebrafish recover an upright posture by the body flexion to the ear-up side.

### Body flexion continues throughout sustained head roll tilt

To further clarify the relationship between the head roll tilt and body flexion, fish’s anterior body was rigidly restrained to a rectangular chamber using an acrylic holder in fish tank water without methylcellulose, and a roll tilt stimulus was applied (Figure 2A). With this setup, fish’s head roll angle was directly coupled with the chamber’s tilt angle throughout the experiment. When a roll tilt stimulus was applied towards the fish’s left-down or right-down side and the tilt was kept for 3 s, fish continued the body flexion to the ear-up side throughout the sustained stimulation (Figure 2B and 2C). In some trials, fish exhibited swim-like lateral body undulation and tail flick to either left or right side during the sustained tilt (Figure S2). As the sustained tilt stimulation was released, the flexed body relaxed. These results further demonstrate the tight correlation between the head roll tilt and body flexion.

**Figure 2.**
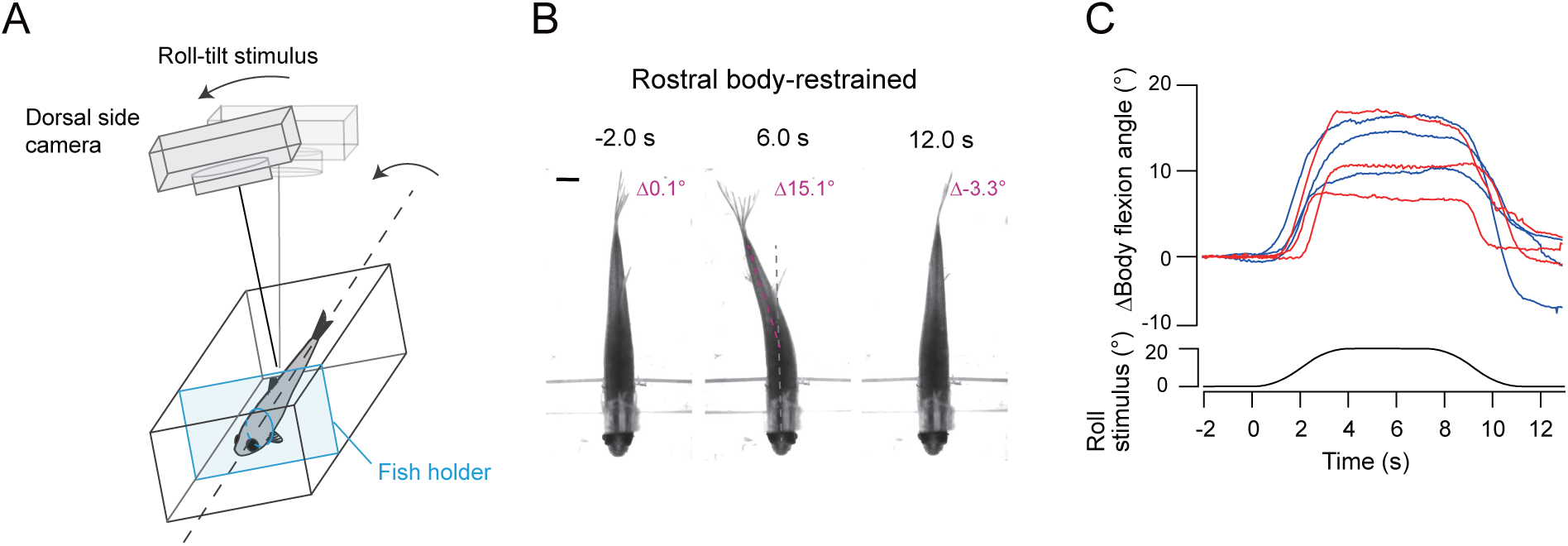
Restrained behavior during static roll tilt. (A) Schematic illustration of restrained behavioral experiments. A fish was restrained and subjected to roll tilt stimuli in a dark room. Its behavior was recorded by a dorsal camera, which tilted together with the fish chamber. (B) Snapshots of a restrained fish during a roll tilt stimulus towards its left-down side. Before stimulus onset, the body is straight (-2.0 s). While the sustained tilt stimulus is applied, the fish maintains body flexion (6.0 s). As the tilt stimulus is released, the fish straightens its body (12.0 s). Scale bars indicate 3 mm. (C) Time courses of body flexion angle in response to roll tilt stimuli. Average traces from six individual fish (5 to 8 trials per fish) are shown (red: *casper* fish; blue: wild-type fish). See also Figure S2.

### Biomechanical model of postural recovery through body flexion

Mechanistically, how does the body flexion contribute to the roll postural recovery? A biomechanical model of the posture control has been proposed in a larval fish study.^12^ The body flexion during roll tilt causes asymmetric positioning of the low-density swim bladder relative to the whole body, resulting in the misalignment of gravity and buoyancy forces, which generates a corrective moment of force for postural recovery.^12^ Building upon this biomechanical framework established in larval zebrafish, we developed a corresponding model for adult zebrafish. In adult zebrafish, the center of mass (COM) and center of volume (COV) are located near the constriction region between the anterior and posterior swim bladder chambers.^11^ Both COM and COV are located on the midline when a fish is at rest: gravity acting on COM and buoyancy acting on COV are balanced (Figure 3 left column). When tilted in the roll axis, the fish flexes its body towards the ear-up side (Figure 3 middle column). The swim bladder shifts toward the ear-down side relative to the whole body, while the head and tail reposition toward the ear-up side. Because the swim bladder contains gas with a density orders of magnitude lower than the rest of the body tissues, the ear-down side positioning of the swim bladder shifts relative positions of COM and COV, which misaligns the gravity and buoyancy forces. This misalignment generates a moment of force that counteracts the roll tilt, thereby restoring an upright posture. According to this model, postural recovery is expected to be disturbed in a swim bladder-deflated fish. Specifically, in the swim bladder-deflated fish, the density of the body is nearly homogeneous. As a result, positions of the COM and COV hardly separate, even when the fish performs body flexion during roll tilt (Figure 3 right column). Consequently, little moment of force is generated, resulting in significantly impaired postural recovery. Additionally, the swim bladder-deflated fish is expected to be prone to roll tilt, since it is not able to efficiently counteract roll tilt through body flexion.

**Figure 3.**
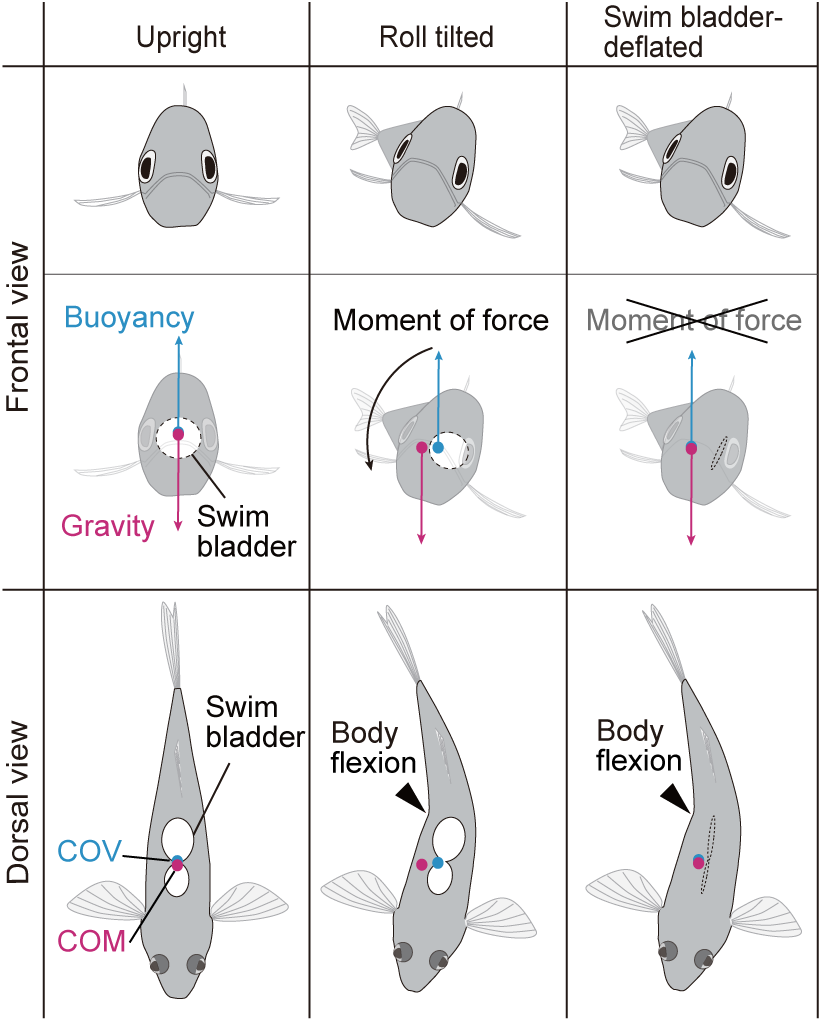
A mechanistic model of static roll posture control through body flexion in adult fish. A biomechanical model of postural recovery from roll tilt by body flexion adapted from a model for larval zebrafish in Sugioka et al.^12^ Gravity and buoyancy forces act at the center of mass (COM) and center of volume (COV), respectively. The COM and COV are located near the constriction region between the anterior and posterior swim bladder chambers.^11^ Left column: At rest, the fish’s COM and COV are located on the midline, and the gravity and buoyancy forces are nearly balanced, enabling the fish to maintain an upright posture. Middle column: When the fish is tilted in the roll direction, it flexes its body towards the ear-up side, causing the swim bladder to shift towards the ear-down side. The extremely low density of the swim bladder results in a shift in the relative positions of the COM and COV. This misalignment of gravity and buoyancy axes generates a moment of force that restores the upright posture. Right column: A fish with a deflated swim bladder flexes its body when tilted in the roll direction. Due to nearly uniform body density, the relative position of COM and COV remains almost unchanged. Consequently, the moment of force is greatly reduced, preventing restoration of upright posture.

### Swim bladder plays a pivotal role in roll posture recovery

To test whether the model holds true in adult fish, as previously demonstrated in larvae, we deflated swim bladder in *casper* fish and performed behavioral analysis (Figure S3). To fairly evaluate the impact of swim bladder deflation, control experiments (sham deflation) were also conducted: this sham procedure involved all the steps performed on the deflated fish with the exception of piercing the swim bladder sac and applying negative pressure to remove the bladder gases. After the sham procedure, fish initiated locomotion within a few minutes, and locomotion speed recovered to levels comparable to pre-procedure levels within the 10-minute recovery period (Figure S4). Furthermore, when roll tilted, sham fish exhibited posture recovery behavior with kinetic parameters similar to those of intact fish (see texts below, Table S1, and STAR Methods for kinetic parameters). These data indicate that the sham procedure including anesthesia did not significantly affect fish behaviors.

In response to roll tilt stimulus, sham fish flexed their body and recovered an upright posture. Figure 4A (top photographs) and Video S2 show examples. When a roll-tilt stimulus was applied towards the fish’s right-down side, a sham fish was slightly tilted to the right-down direction. Simultaneously, the fish flexed its body to the left (ear-up) side. After the stimulus reached its final angle, the head roll angle decreased and the body flexion relaxed to the original state (Figure 4A, right column in the top photographs).

**Figure 4.**
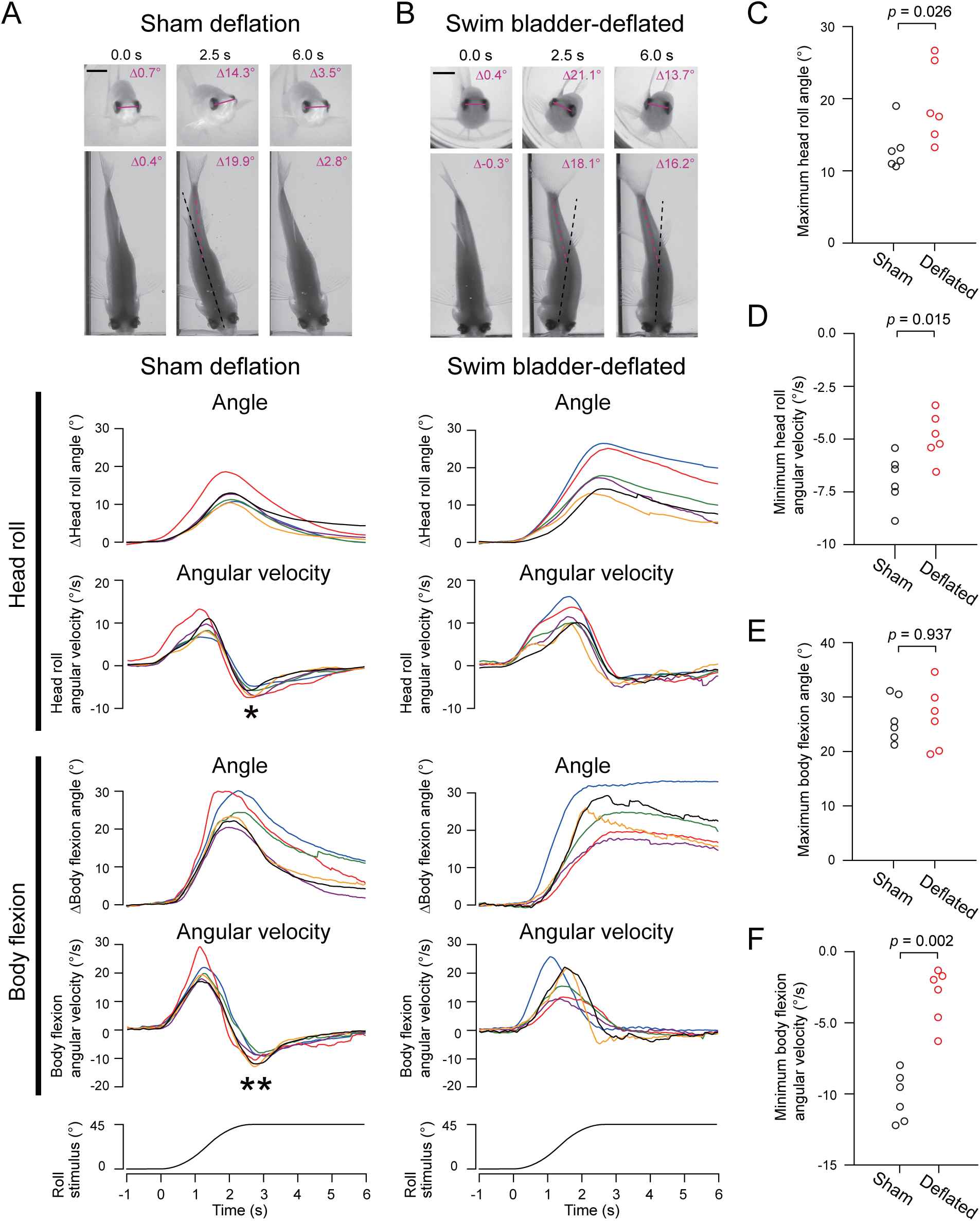
Behavior of static roll posture control in swim bladder-deflated fish. (A and B) Behavioral images and kinetics of sham control and swim bladder-deflated *casper* fish. (Top photographs) Snapshots of a sham fish (A) and swim bladder-deflated fish (B) during a roll tilt stimulus (top row: frontal view; bottom row: dorsal view). Fish tank water contains methylcellulose (A) and methylcellulose and sucrose (B). A sham fish maintains an upright posture before stimulus onset (0.0 s). As the fish is tilted in the right-down direction, it flexes its body towards the left (ear-up) side (2.5 s). The fish recovered upright posture, while its body flexion relaxed to the original state (6.0 s). A deflated fish maintains an upright posture before stimulus onset (0.0 s). As the fish is tilted in the left-down direction, it flexes its body towards the right (ear-up) side (2.5 s). The fish remains roll tilted, while it continues body flexion (6.0 s). Scale bar indicates 3 mm. (Bottom traces) Time courses of changes in head roll angle, head roll angular velocity, changes in body flexion angle, and body flexion angular velocity in response to roll tilt stimuli. Average traces from six individual fish are shown in different colors (4 to 8 trials per fish). In angular velocity traces of head roll and body flexion, there are troughs (asterisk and double asterisks) during postural recovery in sham fish group, while troughs are not clearly observed in deflated fish group. (C-F) Comparison of behavioral parameters between sham group and swim bladder-deflated group. (C) Maximum head roll angle. *p* = 0.026, Mann-Whitney U test. (D) Minimum head roll angular velocity. *p* = 0.015, Mann-Whitney U test. (E) Maximum body flexion angle. *p* = 0.937, Mann-Whitney U test. (F) Minimum body flexion angular velocity. *p* = 0.002, Mann-Whitney U test. See also Figures S3 and S4.

In swim bladder-deflated fish experiments, when a roll-tilt stimulus was applied towards the fish’s left-down side, a fish was slightly tilted to the left-down direction (Figure 4B, top photographs and Video S2). Simultaneously, the fish flexed its body to the right (ear-up) side. In contrast to sham fish, the body flexion did not relax to the original state and the head roll angle only slightly and gradually decreased, after the stimulus reached its final angle (Figure 4B, right column in the top photographs).

To evaluate the effects of swim bladder deflation on behavioral kinetics, angles and angular velocities of the head roll and body flexion were compared (Figure 4A and 4B, bottom traces). In both sham and deflated fish, head roll angle increased as the roll tilt stimulus angle increased. The head roll onset time was not significantly different (0.065 ± 0.052 s in sham; 0.176 ± 0.170 s in deflated, *p* = 0.145, Mann-Whitney U test). Based on our biomechanical model described above, deflated fish were predicted to be more prone to roll tilt, since they cannot efficiently counteract roll tilt through the body flexion. Consistent with this model prediction, deflated fish exhibited larger maximum head roll angles (Figure 4C, *p* = 0.026, Mann-Whitney U test), and showed delayed head roll peak timing (2.024 ± 0.071 s in sham; 2.576 ± 0.152 s in deflated, *p* = 0.002, Mann-Whitney U test) compared to sham controls.

More importantly, swim bladder deflation significantly impaired postural recovery dynamics. While head roll angles gradually decreased in both sham and deflated fish during recovery phase after reaching peak roll angles, the recovery kinetics were slower in deflated fish. This difference was evident in their head angular velocity profiles: during the recovery phase, sham fish showed pronounced troughs in their velocity traces (asterisk in Figure 4A; note that negative velocity reflects rapid recovery movement), whereas such troughs were much less evident in the traces of swim bladder-deflated fish (Figure 4B). Quantitatively, the minimum head roll angular velocity was significantly less negative (indicating slower recovery) in deflated fish compared to sham fish (Figure 4D, *p* = 0.015, Mann-Whitney U test). These data demonstrated that swim bladder plays a critical role in effective postural recovery.

As head roll stimulus angle increased, body flexion angle also increased in both sham and deflated fish. Similarly to the head roll angle changes, the onset time of body flexion was not significantly different (0.238 ± 0.058 s in sham; 0.454 ± 0.208 s in deflated, *p* = 0.093, Mann-Whitney U test), while the peak time was delayed in deflated fish (2.087 ± 0.124 s in sham; 3.304 ± 0.628 s in deflated, *p* = 0.004, Mann-Whitney U test). There was no significant difference in the maximum body flexion angle (Figure 4E, *p* = 0.937, Mann-Whitney U test).

In sham fish, as the stimulus approached the final angle, the body flexion angle began to decrease toward the baseline. In deflated fish, in contrast, the body flexion angle remained high and did not immediately return to the baseline. As described above, sham fish exhibited faster head roll recovery compared to deflated fish. Consequently, the speed of body flexion relaxation was also faster: sham fish exhibited pronounced troughs in their angular velocity traces (double asterisks in Figure 4A; note that negative velocity reflects rapid body relaxation), whereas such troughs were barely detectable in the traces of swim bladder-deflated fish (Figure 4B). The minimum body flexion angular velocity was significantly less negative (indicating slower body relaxation) in the deflated fish group (Figure 4F, *p* = 0.002, Mann-Whitney U test). These results demonstrated that deflating the swim bladder disturbs posture recovery despite fish performing sustained body flexion during roll tilt. Collectively, these findings suggest that adult zebrafish, similar to larvae, employ body flexion for static roll posture control, where the swim bladder plays a crucial role in effective posture recovery.

### Body flexion with intact swim bladder generates recovery moment of force

To examine whether body flexion with intact swim bladders generates moment of force, we analyzed passive movement of chemically fixed, body-flexed fish specimens before and after swim bladder deflation. Fish were euthanized and chemically fixed in a body-flexed posture (Figure 5A, top). The body flexion angles (θ1 in Figure 5A) were between 14 and 24 degrees in the 4 specimens examined. The fixed fish specimen was held by a pair of forceps in a chamber filled with fish tank water. To mimic the body orientation of live fish during roll tilt (Figure 1 and 4), the fixed specimen was held in a roll-tilted orientation with its body flexed to the ear-up side (Figure 5A, bottom). Care was taken to apply equivalent roll tilt angles (θ2 in Figure 5A) across before-deflation and after-deflation conditions (before deflation: 24.3 ± 4.7°; after deflation: 24.0 ± 5.5°). The tilted specimen was gently released and passive movement was recorded by a front camera.

**Figure 5.**
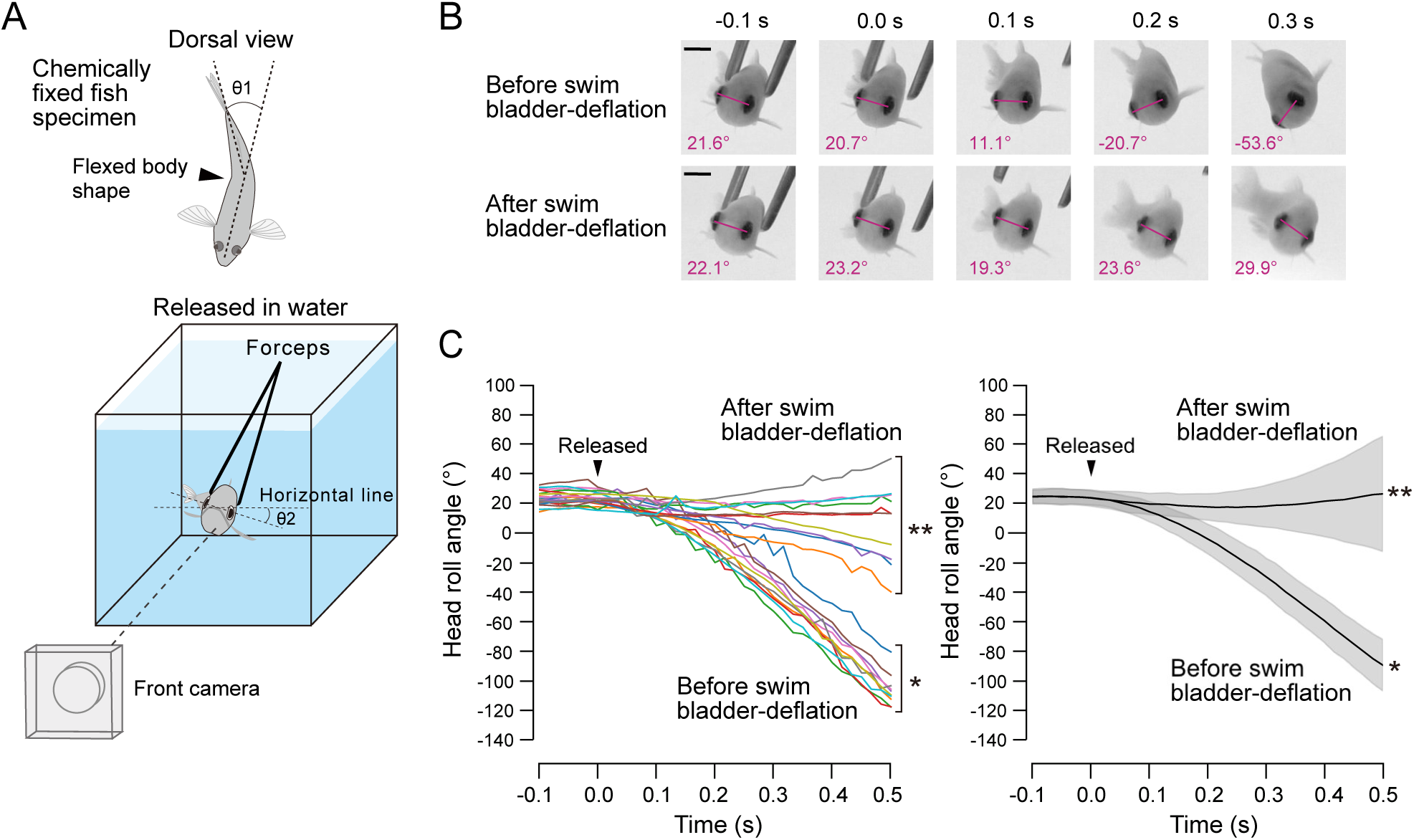
Fixed-specimen passive movement experiments. (A) Schematic illustration of fixed-specimen passive movement experiments. A fish was euthanized and chemically fixed in a body-flexed posture with body flexion angle (θ1) approximately 20 degrees. The fixed fish specimen was held by a pair of forceps in a chamber filled with fish tank water. The held specimen was manually roll tilted approximately 24 degrees (θ2) with its body flexed to the ear-up side. The tilted specimen was released and passive movement was recorded by a front camera. (B) Snapshots of passive movement of a released fish specimen before (top row) and after (bottom row) swim bladder deflation. Before deflation, the specimen rolls toward ear-up side and continues rolling. After deflation, the specimen does not roll. Scale bar indicates 3 mm. (C) Time courses of head roll angle before (asterisk) and after swim bladder deflation (double asterisks). (Left) Ten trials in each condition from an example fish. (Right) Data from 4 fish. Black traces represent mean and gray areas represent standard deviation.

Once the constraint by the forceps was removed, the specimen before swim bladder deflation started passive rolling toward the ear-up side (Figure 5B, top photographs and Video S3). This movement was consistent with the biomechanical model (Figure 3) and was similar to postural recovery of intact and sham fish (Figures 1 and 4A). A key difference between intact/sham fish and chemically fixed specimens was the presence or absence of body straightening during postural recovery. In intact and sham fish, the body returns to the symmetric, straight shape as the head roll angle approaches zero (Figure 1B and 1C), which erases recovery moment of force and allows fish to settle into an upright roll posture. In chemically fixed specimens with body-flexed shape, the body flexion angle remains constant regardless of head roll angle, which likely generates moment of force that keeps the specimen rolling passing through upright orientation. As expected, the fixed specimens continued rolling even after they reached an upright orientation, and head roll angle continuously decreased (asterisk in Figure 5C, left panel). Similar movements were observed in all 4 fish specimens (asterisk in Figure 5C, right panel).

To examine whether swim bladder deflation disturbs the passive roll recovery, after the passive movement experiments described above, the swim bladder was deflated and the specimens were again subjected to the same passive movement experiments. Deflated specimens did not consistently roll toward the ear-up direction (Figure 5B, bottom photographs and Video S3): they did not roll or slightly rolled in a random direction (double asterisks in Figure 5C, left panel). Similar movements were observed in all 4 fish specimens (double asterisks in Figure 5C, right panel). Head roll angular velocities were calculated from the period from 0.2 to 0.5 s after the release. Angular velocity was significantly reduced after swim bladder deflation (before deflation:-290.0 ± 48.4°/s; after deflation: 31.0 ± 111.2°/s; *p* = 1.3e-22, Welch’s t-test). These results further demonstrated that body flexion with intact swim bladders is sufficient to produce passive recovery moment of force for roll postural recovery.

## Discussion

The present study has established an experimental paradigm for analyzing roll postural control behavior in adult zebrafish and demonstrated that when slightly roll tilted, adult zebrafish exhibit body flexion to recover an upright posture (Figure 1). Both the tight correlation between body flexion and head roll angle (Figure 2) and the dependence on swim bladder for postural recovery (Figure 4) suggest that the mechanisms underlying roll postural maintenance through body flexion are similar to those in larvae (Figure 3).^12^ The passive movement experiment using chemically fixed fish specimens demonstrated that the flexed body with intact swim bladder generates recovery moment of force (Figure 5). The conservation of these mechanisms between larvae and adults suggests that body flexion serves as a fundamental strategy for static roll postural maintenance throughout their life.

Previous researches on larvae suggested that the otolith organs in the vestibular system sense roll tilt and induce the body flexion. ^12,13,15,16^ Not only the otolith system, but fish may also use sensory information from eyes, semicircular canals, and lateral line, to perform the body flexion during roll tilt. In our experiments, however, the body flexion was observed in dark conditions. Therefore, visual inputs are not necessary for the body flexion. Moreover, in restrained behavioral experiments, fish maintained the body flexion throughout the sustained tilt stimulus, during which angular velocity and water flow were minimal, while gravitational input from otolith organs persisted. This observation suggests that the gravity-sensing otolith organs play a primary role in inducing this behavior.

Neural circuits and muscles that induces the body flexion behavior during roll tilt were identified in zebrafish larvae.^12,13,16,17,26,27^ A vestibular nucleus (tangential nucleus) receives head roll tilt signals from vestibular primary afferents and send neural signals to a group of reticulospinal neurons in the nucleus of the medial longitudinal fasciculus that projects to the spinal cord.^12^ These neuronal nuclei also exist in adult fish.^28–30^ This conservation of the neuronal nuclei and similarity of the body flexion behavior between larvae and adults suggest that the same neural circuits likely play a principal role in inducing the body flexion in adult fish. Regarding the muscles responsible for the body flexion during roll tilt, adult zebrafish appeared to flex their body near the middle of the body where the caudal chamber of the swim bladder is located (Figure 1B and 2B). In larvae, muscle contraction during roll tilt also appears to occur only near the swim bladder.^12^ Consistent with this, the PHM muscles lateral to the swim bladder generate body flexion, with slow-type PHM primarily responsible for this movement.^12,17^ These data suggest that the slow-type PHMs also generate the body flexion in adults. Muscular organization in adult fish, however, is more complex and extensively developed compared to larvae.^18,20,31,32^ Thus, other muscles, such as trunk slow muscles that are recruited during weak and sustained body movements, may also contribute to the body flexion.^33^ Experimentally examining the causal relationship between the body flexion and neuromuscular cell populations in adult fish is technically challenging due to their size, opaque tissues, ossified skull, and lack of gene promoters for selectively targeting a specific cell population. However, future advancements in technology, including optical, genetic, and chemical approaches, as well as combination of these, may help overcome these limitations and clarify neural and muscular mechanisms.

In a biomechanical model, the body flexion toward the ear-up side displaces the low-density swim bladder toward the ear-down side, resulting in separation of the COM and COV (Figure 3). The magnitude of the recovery moment of force is proportional to the distance between the COM and COV, which is maximized under a condition where the swim bladder, the least dense part of the body, is located near the COM, and the body flexes at the position near the swim bladder. The body organization and the body flexion in larval zebrafish meet this condition, indicating that the body flexion is a reasonable, energy-efficient behavior for postural correction.^12^ Similarly in adult zebrafish, the COM—positioned at approximately one third of the total body from the rostral end—is located around a constriction region of the swim bladder separating the rostral and caudal chambers.^11^ Regarding the body flexion region, as discussed above, the body flexed around the middle of the body (Figures 1B, 2B, 4A and 4B), the location of which was slightly caudal to the swim bladder. Thus, the flexion region seems to slightly caudally deviate from the most biomechanically efficient location. This subtle deviation may be attributed to the musculoskeletal organization in adult fish. In the abdominal region in the rostral body, bony structures increase the body stiffness: lateral angular stiffness of the vertebral column is higher in the abdominal region than the caudal body region,^34^ and the rib cage bones covering internal organs presumably increase the stability in the abdominal region. These morphological characteristics make the rostral body flexion challenging and constrain the flexion region primarily to the middle and caudal body.^35,36^ Considering these morphological constraints, the flexion at the middle of the body is presumably an adaptive compromise between biomechanical efficiency and structural limitations. While the results from the swim bladder deflation experiments in live fish (Figure 4) and fixed fish specimens (Figure 5) supported the biomechanical model (Figure 3), the precise locations of the COM and COV during body flexion have not been provided. Further investigation through morphometric analyses would help address these questions.

Swim bladder-deflated fish were prone to roll tilt, and their maximum head roll angle was larger than sham control (Figure 4C). Previous studies showed that larval zebrafish increase body flexion angle in response to increasing vestibular stimulus mimicking head roll.^15,16^ Given that larval and adult zebrafish employ similar roll posture control mechanisms, deflated adult zebrafish in the present study would attempt to counteract roll tilt by increasing their body flexion angle. However, the maximum body flexion angle was comparable between control and deflated groups, despite deflated fish experiencing greater postural perturbations (Figure 4E). These data suggest that adult fish face morphological constraints that limit their compensatory body flexion capacity. Unlike flexible larvae, adult fish possess increased body stiffness due to ossified skeletal structures. This may restrict the range of achievable body flexion angles in adult fish.

Previous studies showed that during tilt in the roll axis, fish exhibit body undulation with asymmetric motor pattern that recovers their posture.^4,5,8^ Consistently, in the present study, swim-like vigorous body movement and fin beating were observed when adult zebrafish was roll tilted. Intriguingly, in experiments where the fish’s rostral body was restrained and sustained tilt stimulation was delivered, fish occasionally exhibited a swim-like body undulation and tail flick behaviors in addition to their body flexion as larvae do (Figure S2).^13,15,16^ This indicates that fish maintain roll posture primarily through static control via body flexion, and recruit dynamic control through vigorous body movements when static control alone is insufficient.

What are the advantages of static posture control through body flexion? Compared to swimming and fin movements, maintaining posture through slight body flexion is less visually conspicuous and therefore likely reduces the risk of being detected by predators.^37,38^ Additionally, body flexion presumably requires less energy compared to vigorous axial undulation during swimming.^39^ This is particularly beneficial during rest periods when maintaining postural stability while conserving energy is crucial.

The mechanism of posture control by body flexion is likely a widely used strategy employed by many aquatic organisms with low-density organs. While our study focused on the swim bladder, not only the swim bladder in many fish species, but also lungs in lungfish and lipid-rich liver in sharks may play a crucial role when these animals flex their body upon roll tilt.^40–42^ Further investigation across a broader range of species will elucidate the generality of the body flexion-mediated, static roll posture control mechanism.

### Limitations of the study

Swim bladder-deflated fish showed slow but noticeable posture recovery (Figure 4). Technical limitations in the deflation procedure might explain this slow recovery: the swim bladder may not have been fully deflated. While the posterior chamber of the swim bladder was deflated, the anterior chamber was difficult to observe and therefore its deflation was not confirmed. Small amount of residual gas might have generated a small recovery moment of force during body flexion. In the fixed specimen experiments, since specimen viability was not a concern, we were able to apply stronger negative pressure to more extensively remove gas from the swim bladder. This might have produced more pronounced impairment of posture recovery (Figure 5).

Most behavioral experiments were conducted in viscous water conditions using methylcellulose in order to efficiently deliver roll tilt stimulus and clearly observe posture control responses. Due to the increased viscosity, the behavioral kinematic profiles measured under viscous water conditions were likely slowed relative to those under normal water conditions. The viscous medium may also affect the relative contributions of static versus dynamic postural control strategies, as the increased drag forces could favor sustained postural adjustments over rapid swimming movements. However, fish exhibited body flexion during roll tilt under normal water condition in our restrained experiments (Figure 2), and experiments using chemically fixed specimens demonstrated that the fundamental body flexion mechanism operates effectively in normal water, suggesting that while the temporal dynamics and the relative contributions of static versus dynamic postural control strategies may be altered by viscosity, the underlying biomechanical principles remain valid.

Although the present study has demonstrated that body flexion plays a crucial role in roll posture control, this mechanism represents only one component of the complex postural control repertoire that fish employ to maintain an upright posture. Previous studies on fish postural control have primarily focused on dynamic control mechanisms. Fin movements vary considerably among fish species. Some species skillfully control their fins to maintain posture even while they do not locomote.^43–45^ In these species, fin movements, particularly the pectoral fins, may play a primary role in maintaining posture. The present study focused on static posture control and did not provide quantitative analysis of the relative contributions and frequencies of static versus dynamic control strategies. As discussed above, some fish species may predominantly use dynamic posture control. Given that fish species have evolved diverse survival strategies across various aquatic environments, future comparative studies examining how postural control strategies vary among species with different body morphologies and ecological niches would provide important insights into the adaptive significance of static versus dynamic postural control mechanisms.

## Resource availability

### Lead contact

Further information and requests for resources and reagents should be directed to and will be fulfilled by the Lead Contact, Masashi Tanimoto.

## Materials availability

This study did not generate new unique reagents.

## Data and code availability

All data reported in this paper will be shared by the lead contact upon request. All original code has been deposited at Zenodo at https://zenodo.org/records/14551001 and is publicly available as of the date of publication. Any additional information is available from the lead contact upon request.

## Supporting information

Fugure S1

Fugure S2

Fugure S3

Fugure S4

## Acknowledgements

The authors are grateful to Higashijima lab members for their help with fish care and discussion. This work is supported by Japan Society for the Promotion of Science KAKENHI Grant Numbers JP20K06866 and JP23K05983.

## Author contributions

R.N., T.K., S.H., and M.T. designed the research; T.K. built a prototype of the experimental apparatus under the supervision of M.T.; R.N. established the apparatus, performed experiments, and analyzed data under the supervision of M.T.; R.N. and M.T. wrote the manuscript.

## STAR Methods

Key resources table

## EXPERIMENTAL MODEL AND STUDY PARTICIPANT DETAILS

### Fish

Adult wild-type and *casper*^25^ zebrafish (38-45 mm in total length) were maintained under light-dark cycles (10:14 or 12:12 hours) under standard conditions.^46^ For all experiments, both sexes of fish that displayed normal body morphology and maintained a dorsal-up posture while at rest were used. For unrestrained behavioral experiments of intact fish, 2 males and 2 females of *casper* fish and 2 males and 2 females of wild-type fish were used. For restrained behavioral experiments, 2 males and 1 female of wild-type fish and 1 male and 2 females of *casper* fish were used. For unrestrained behavioral experiments of *casper* fish with swim bladder deflation, 3 males and 3 females were used for sham deflation, and 3 males and 3 females were used for deflation. For locomotion experiments, 1 male and 3 females of *casper* fish were used. For fixed specimen passive movement experiments, 2 males and 2 females of *casper* fish were used. All experiments were conducted according to animal experiment protocols approved by the animal care and use committees of the National Institutes of Natural Sciences.

## METHODS DETAILS

### Unrestrained behavioral experiments

The fish chamber consisted of two acrylic cylinders of different sizes (22 mm [inner diameter] × 50 mm [length] × 1.5 mm [thickness] and 46 mm [inner diameter] × 10 mm [length] × 2 mm [thickness]) connected by an acrylic ring plate. The two cylinders were separated by a transparent, fluorinated ethylene propylene sheet (NR0538-002, Fronchemical), which had similar refractive index as water. Small holes were made on this sheet so that water passes between the two cylinders. Both ends of the connected chamber were covered with acrylic lids. The two cylinders served different purposes: the smaller cylinder housed a fish, while the larger cylinder supplied oxygen and, by manually tilting the chamber, removed small air bubbles from the smaller cylinder. Air bubble was maintained in the larger cylinder for oxygen supply. Since it was difficult to consistently tilt adult fish and reliably observe their postural responses in normal water conditions, presumably due to their large inertia and rapid postural recovery, behavioral experiments were conducted under viscous water conditions. The chamber was filled with fish tank water containing 0.8% methylcellulose (22223-65, Nacalai Tesque, inc.), and a fish was placed in the smaller cylinder.

The rotation stimulus device was composed of a motorized rotation stage (DDR100/M, ThorLabs) and supporting mechanical components. The fish chamber was mounted on this device. The stage was driven by a stage controller (BBD301, ThorLabs) using a software (Kinesis version 1.14.49, ThorLabs). Fish were illuminated by an infrared light-emitting diode (IR-LED) plate (LFL-100IR2-850, CCS) from the ventral side and another IR-LED (HLV3-22IR860, CCS) from the caudal side. Experiments were conducted in a dark room to eliminate visual cues. The chamber was rotated 45° in the roll direction (maximum velocity: 40°/s, acceleration and deceleration: 25°/s²) unless otherwise noted. Fish behavior was recorded at 60 frames per s (fps) by two cameras (HC-VX992MS, Panasonic) from the frontal and dorsal sides. The timing of rotation stimulus and camera image frames were synchronized by a flash of an IR-LED, which was oriented towards two cameras and distinct from the illumination IR-LEDs. In each trial, a voltage pulse was sent from the rotation stage controller to the IR-LED through a pulse generator (SEN7203, Nihon Kohden) at 1 s before the rotation started.

Since swim bladder-deflated fish were prone to roll tilting, they exhibited dynamic posture control through swimming and tail flick more frequently compared to intact and sham fish when subjected to the standard roll stimulation described above. To better observe static posture control via slight body flexion, a weaker rotation stimulus of 35-45° (maximum velocity: 30-40°/s, acceleration and deceleration: 20-25°/s²) was applied. The reduced buoyancy caused by swim bladder deflation was compensated by 5-32% (in most cases 10-25%) sucrose (196-00015, Fujifilm Wako), depending on neutral buoyancy of individual fish, in fish tank water containing 0.8% methylcellulose (22223-65, Nacalai Tesque, inc.) solution. For wild-type fish, which were slightly larger than *casper* fish, a cylinder chamber with larger dimensions (26 mm [inner diameter] × 50 mm [length] × 2 mm [thickness] and 46 mm [inner diameter] × 10 mm [length] × 2 mm [thickness]) was used.

### Swim bladder deflation

Pigment-less *casper* fish were used for easier identification of the swim bladder position. Fish were anesthetized with 0.01% MS-222 (Ethyl 3-aminobenzoate methanesulfonate, A5040, Sigma-Aldrich) for 10 minutes, and after confirming the absence of response to tactile stimuli, a syringe needle (27G [NN-2719S, Terumo] or 30G [Dentronics]) was gently inserted into the caudal chamber of the swim bladder from a lateral and slightly dorsal and caudal direction. Negative pressure was gently and gradually applied to the syringe to remove the gas from the swim bladder. Successful swim bladder deflation was confirmed by the caudal chamber of the swim bladder becoming invisible. Following deflation, fish were returned to tank water, allowed to rest for 10 minutes before behavioral experiments.

### Sham deflation

Sham swim bladder deflation was performed almost exactly as those for deflation procedure described above. The only modification was that the swim bladder sac was not penetrated by a syringe needle and swim bladder gas was not removed for sham deflation. Even slight piercing of the swim bladder sac with a syringe needle resulted in gas leakage, which would have compromised the intended control condition. Therefore, sham fish underwent identical handling, surgical preparation, and recovery procedure to those used for deflated fish, but with the swim bladder left intact avoiding penetration of the swim bladder sac.

### Locomotion before and after sham deflation procedure

*casper* fish were used. Individual fish was placed in a tank (260 mm [length] × 150 mm [width] × 175 mm [height]) filled with fish tank water (depth: 70 mm), and behavior was recorded at 60 fps for 11 minutes from the top by a camera (HC-VX992MS, Panasonic). Sham swim bladder deflation was performed as described above. After the sham procedure, fish was again placed in the tank and behavior was recorded.

### Restrained behavioral experiments

Fish were anesthetized with 0.01% MS-222 for 10 minutes. The fish’s body between the operculum and pectoral fin was immobilized by a separable pair of acrylic plates that have a ditch: an elliptical hole (major axis: 6.0-6.75 mm; minor axis: 4-5 mm) was formed when these plates were combined together. To stabilize the body position, a small amount of flexible silicone (Macks Earplugs) was applied to the space between the hole and the fish body. The plate holding the fish was then placed into a groove of an acrylic chamber (60 mm [length] × 30 mm [width] × 45 mm [height]) filled with fish tank water without methylcellulose. An open-topped rectangular box (55 mm [length] × 30 mm [width] × 20 mm [height]) was placed over the fish to prevent image distortion when tilt stimulus was applied and the water surface tilted. Fish were allowed to rest for at least a few minutes before stimulus delivery and camera recording began. Fish was illuminated by an IR-LED (LFL-100IR2-850, CCS) placed below the chamber. The rotation stimulus device was the same as that described in ‘Unrestrained behavioral experiments’ section. The chamber was mounted on the rotation stimulus device, and 20° roll-tilt stimulation (maximum velocity: 10°/s, acceleration and deceleration: 5°/s²) was applied and held for 3 s before returning to its original position. Fish behavior was recorded at 25 fps by a camera (daA720-520um, Basler) with a lens (13VM308ASIRII, Tamron) from the dorsal side using software (pylon Viewer version 6.2.0.8205, Basler). The camera tilted together with the fish chamber. The timing of the rotation stimulus and the camera image frames were synchronized by voltage pulses. Specifically, from 2 s before the rotation started in each trial, image frame acquisition was triggered by voltage pulses from the rotation stage controller through a pulse generator (SEN-7203, Nihon Kohden).

### Fixed-specimen passive movement experiments

Fish were euthanized by placing them with 0.04% MS-222 for 30 minutes. The fish specimens were restrained on a sponge plate using metal pins, with the body maintained in a bent posture such that the body flexion angle was approximately 20 degrees. The body bend direction was to the left in 2 fish and to the right in 2 fish. The fish specimens were fixed with 4% paraformaldehyde (158127, Sigma-Aldrich) in 1x phosphate buffered saline (PBS) for 30 minutes at room temperature. After brief rinse with 1x PBS, the fish specimens were photographed from the dorsal direction by a camera (HC-VX992MS, Panasonic). The body flexion angle of the fixed fish specimens was between 14 and 24 degrees in 4 specimens examined.

The fixed fish specimen was held by a pair of forceps in a chamber filled with fish tank water. The held specimen was manually roll tilted from the dorsal-up orientation with its body flexed to the ear-up side. The tilted specimen was released and passive movement was recorded by a front camera (HC-VX992MS, Panasonic) at 60 fps. Care was taken to minimize external forces to the specimen after the release. Swim bladder deflation was performed as described above. Since specimen viability was not a concern, swim bladder deflation was performed more extensively than in live fish, allowing for stronger negative pressure application and more complete gas removal.

## QUANTIFICATION AND STATISTICAL ANALYSIS

### Image analysis of live fish experiments

Captured videos were analyzed with a custom program written in Python (version 3.12.7). The data analysis employed Python libraries including NumPy, OpenCV, Matplotlib, pandas and scikit-image^47–51^. Videos from unrestrained behavioral experiments were downsampled to 30 fps, which was sufficient to capture behavioral dynamics. Videos from restrained behavioral experiments were analyzed at the original frame rate. For analysis, videos were converted into series of TIFF images using OpenCV. The head roll angle was calculated as follows: each image from the frontal camera was binarized to isolate pixels containing the eyes, and the xy coordinates of each eye’s center were estimated. The head roll angle was then measured as the angle between the line connecting the centers of the two eyes and the horizontal line.

The body flexion angle was calculated using images from the dorsal camera. Each image was binarized to extract pixels containing the fish while excluding the fins. The fish’s midline, from snout to tail, was divided into 23 segments. The body flexion angle was determined by calculating the angle between two lines: (1) the line connecting the second and twelfth points from the tail, and (2) the line connecting the twelfth and twenty-second points. These points were selected since the two lines best represent the body flexion in most cases. In image frames where the fish’s snout or tail overlapped with the contour of the cylinder in the images, the point closest to the body flexion point (either the thirteenth, fourteenth, or fifteenth point) was chosen as the middle point. The angle was calculated between (1) the line connecting the second and middle points, and (2) the line connecting the middle and twenty-second points. The dorsal camera images were distorted by optical refraction through the curved surface of the acrylic cylinder, resulting in slight expansion of image objects in the cylinder’s short-axis direction. The magnitude of distortion increased from the top to the bottom of the cylinder along the depth axis. It also increased from the center to the edge along the cylinder’s short-axis direction in horizontal planes, except in the middle horizontal planes where the image slightly shrank near the cylinder’s edge. However, since the body flexion angle was measured from the body midline, this image distortion would minimally affect the angle measurements. Also, since the fish’s location was randomly distributed in the cylinder across multiple trials, any potential effects of image distortion would be averaged out, thus having minimal impact on the angle analysis. Therefore, the body flexion angle was calculated from raw images.

For all measurements except dynamic movement analysis (Figure S2), image frames where fish exhibited swimming or tail flick were excluded from the dataset. Supplementary videos were made from image series using a software (Clipchamp, Microsoft).

### Image analysis of locomotion experiments

Captured videos were analyzed with a custom program written in MATLAB (version R2023a). In each image frame, background was subtracted, binarized to find pixels containing fish, and the centroid position was estimated. Instantaneous locomotion speed was calculated from the centroid position changes and smoothed by 60-frame moving average. Locomotion speed was then calculated as the mean instantaneous speed per minute.

### Image analysis of fixed-specimen passive movement experiments

Head roll angle was calculated from captured videos using the same methods described above for live fish. Videos were analyzed at the original frame rate. For image frames where the custom tracking program failed to detect eye positions, these eye positions were manually annotated. Only trials where fish specimens showed no visible influence of external forces from the pair of forceps after release were included in the analysis. For each individual fish specimen, 10 trials each were randomly selected from the before-deflation and after-deflation conditions.

### Definition of Measured Values and Statistical Analysis

Data was analyzed using a custom code written in Python (version 3.12.7). A single trial where head roll angle exceeded more than 40 degrees was excluded from the analysis. To extract slow changes in head angle, a 21-frame moving average was applied to raw head angle data. Raw body flexion angle data was smooth enough, and therefore raw angle data was used. All time-series head roll and body flexion angle data were offset by subtracting the pre-stimulus baseline that was average angle during the initial 1 s of each trial. The maximum head roll angle and body flexion angle were defined as the peak values after the stimulus onset. Head roll angular velocity was calculated from the smoothened angle data in each trial using the numpy.gradient function from the Python NumPy library, and a 15-frame moving average was applied. For body flexion angular velocity, a 25-frame moving average was applied to raw angle data. Angular velocity was then calculated from the smoothened angle data in each trial using the numpy.gradient function, and a 13-frame moving average was applied. The minimum angular velocity was defined as the lowest value after the tilt stimulus onset. Onset time for head roll and body flexion angles were defined as the first time point at which the angles after the stimulus onset exceeded 2× standard deviation during 1 s pre-stimulus period. Peak time for head roll and body flexion angles were defined as the time point at which the angles after stimulus onset reached their maximum value. For fixed-specimen passive movement, head roll angular velocities were calculated as the slope of linear regression fitted to angle data in each trial during the period from 0.2 to 0.5 s after release. The effect of swim bladder deflation on posture recovery was evaluated by testing angular velocity values using the two-sided Welch’s t-test. All statistical analyses were performed using the SciPy tool (version 1.13.1)^52^. Except the fixed-specimen passive movement analysis, the mean value for each individual was calculated from in the recorded trials. Using the individual means, the overall mean and standard deviation were calculated for intact, sham, and swim bladder-deflated fish. The effect of swim bladder deflation on posture recovery was evaluated using the two-sided Mann-Whitney U test.

## Supplemental information

Figure S1. Unrestrained behavioral experiments in wild-type fish, related to Figure 1.

Figure S2. Dynamic movement during roll tilt, related to Figure 2.

Figure S3. Swim bladder deflation, related to Figure 4.

Figure S4. Locomotion before and after sham deflation procedure, related to Figure 4.

Table S1. Kinematic parameters in intact, sham, and deflated fish, related to Figure 4. Video S1. Static posture control by body flexion during roll-tilt stimulus in intact adult *casper* zebrafish, related to Figure 1

Video S2. Static posture control by body flexion during roll-tilt stimulus in sham deflation and swim bladder-deflated adult *casper* zebrafish, related to Figure 4

Video S3. Passive movement of fixed fish specimens before and after swim bladder deflation, related to Figure 5

Figure S1. Unrestrained behavioral experiments in wild-type fish.

(A, B) Unrestrained behavioral experiments in wild-type adult zebrafish performed in a dark room. (A) Snapshots of a fish during a roll tilt stimulus towards its left-down side (top row: frontal view; bottom row: dorsal view). Magenta lines connecting the center of eyes show head roll angle. Black and magenta dashed lines denote the midlines of the rostral and caudal body, respectively. Changes in the head roll angle and body flexion angle relative to the pre-stimulus baselines are indicated at the upper right. Before stimulus onset, the fish maintains an upright posture (0.0 s). As the fish is tilted in the roll direction, it flexes its body towards the ear-up side (2.5 s). The fish recovers its upright posture as it straightens its body (5.0 s). Scale bars indicate 3 mm. (B) Time courses of changes in head roll angle and body flexion angle in response to roll tilt stimuli. Average traces from four individual fish are shown in different colors (6 to 9 trials per fish).

Figure S2. Dynamic movement during roll tilt.

Time course of body flexion angle in response to sustained roll tilt stimulation in body-restrained experiments. Example 5 trials where fish exhibited dynamic movement are shown in different colors.

Figure S3. Swim bladder deflation.

Lateral view images of pigment-less *casper* fish before and after swim bladder deflation. Scale bar indicates 5 mm.

Figure S4. Locomotion before and after sham deflation procedure.

Average locomotion speed per minute before and after sham deflation procedure. Locomotion speed recovered to levels comparable to pre-procedure levels in a few minutes.

Table S1. Kinematic profiles of head roll and body flexion in intact, sham, and deflated fish.

Kinematic parameters in intact, sham, and deflated fish (mean ± standard deviation). N indicates number of fish. Mann-Whitney U test was used for statistic comparisons. U denotes the test statistic. Asterisks indicate *p* values smaller than 0.05.

Video S1. Static posture control by body flexion during roll-tilt stimulus in intact adult zebrafish.

Front and dorsal view video clips in intact *casper* fish. Video plays at real-time speed.

Video S2. Static posture control by body flexion during roll-tilt stimulus in sham deflation and swim bladder-deflated adult zebrafish.

Front and dorsal view video clips in sham deflation (first half) and swim bladder-deflated (second half) *casper* fish. Video plays at real-time speed.

Video S3. Passive movement of fixed fish specimens.

Front view video clips of a fixed fish specimen before (first half) and after (second half) swim bladder deflation. Video plays at ×0.25 speed.

## References

1. Horak, F. B. Postural orientation and equilibrium: what do we need to know about neural control of balance to prevent falls? Age Ageing 35, ii7–ii11 (2006).

2. Le Mouel, C. & Brette, R. Mobility as the Purpose of Postural Control. Front Comput Neurosci 11, (2017).

3. Maki, B. E., Mcilroy, W. E. & Fernie, G. R. Change-in-support reactions for balance recovery. IEEE Engineering in Medicine and Biology Magazine 22, 20– 26 (2003).

4. Zelenin, P. V., Orlovsky, G. N. & Deliagina, T. G. Sensory-Motor Transformation by Individual Command Neurons. The Journal of Neuroscience 27, 1024–1032 (2007).

5. Bagnall, M. W. & McLean, D. L. Modular Organization of Axial Microcircuits in Zebrafish. Science (1979) 343, 197–200 (2014).

6. Webb, P. W. & Weihs, D. Hydrostatic stability of fish with swim bladders: not all fish are unstable. Can J Zool 72, 1149–1154 (1994).

7. Ehrlich, D. E. & Schoppik, D. Control of Movement Initiation Underlies the Development of Balance. Current Biology 27, 334–344 (2017).

8. Zelenin, P. V., Grillner, S., Orlovsky, G. N. & Deliagina, T. G. The pattern of motor coordination underlying the roll in the lamprey. Journal of Experimental Biology 206, 2557–2566 (2003).

9. Fath, M. & Tytell, E. D. Day and night posture of the bluegill sunfish (Lepomis macrochirus). Preprint at 10.1101/2023.07.13.548884 (2023).

10. Webb, P. W. & Weihs, D. Stability versus Maneuvering: Challenges for Stability during Swimming by Fishes. Integr Comp Biol 55, 753–764 (2015).

11. Robertson, G. N., Lindsey, B. W., Dumbarton, T. C., Croll, R. P. & Smith, F. M. The contribution of the swimbladder to buoyancy in the adult zebrafish (*Danio rerio*): A morphometric analysis. J Morphol 269, 666–673 (2008).

12. Sugioka, T., Tanimoto, M. & Higashijima, S. Biomechanics and neural circuits for vestibular-induced fine postural control in larval zebrafish. Nat Commun 14, 1217 (2023).

13. Beiza-Canelo, N. et al. Magnetic actuation of otoliths allows behavioral and brain-wide neuronal exploration of vestibulo-motor processing in larval zebrafish. Current Biology 33, 2438–2448.e6 (2023).

14. Migault, G. et al. Whole-Brain Calcium Imaging during Physiological Vestibular Stimulation in Larval Zebrafish. Current Biology 28, 3723–3735.e6 (2018).

15. Favre-Bulle, I. A., Stilgoe, A. B., Rubinsztein-Dunlop, H. & Scott, E. K. Optical trapping of otoliths drives vestibular behaviours in larval zebrafish. Nat Commun 8, 630 (2017).

16. Natalia Beiza Canelo. Postural control in larval zebrafish: vestibular behavior repertoire and a new method to study it. Portail HAL Sorbonne Université (2022).

17. Thiele, T. R., Donovan, J. C. & Baier, H. Descending Control of Swim Posture by a Midbrain Nucleus in Zebrafish. Neuron 83, 679–691 (2014).

18. Thorsen, D. H. & Hale, M. E. Development of zebrafish (Danio rerio) pectoral fin musculature. J Morphol 266, 241–255 (2005).

19. Ma, X. & Xu, X. A Swimming-based Assay to Determine the Exercise Capacity of Adult Zebrafish Cardiomyopathy Models. Bio Protoc 11, (2021).

20. McMenamin, S. K. & Parichy, D. M. Metamorphosis in Teleosts. in Current Topics in Developmental Biology vol. 103 127–165 (Academic Press Inc., 2013).

21. McHenry, M. J. & Lauder, G. V. Ontogeny of form and function: Locomotor morphology and drag in zebrafish (Danio rerio). J Morphol 267, 1099–1109 (2006).

22. Burris, B., Jensen, N. & Mokalled, M. H. Assessment of Swim Endurance and Swim Behavior in Adult Zebrafish. Journal of Visualized Experiments (2021) doi:10.3791/63240.

23. Mauguit, Q., Olivier, D., Vandewalle, N. & Vandewalle, P. Ontogeny of swimming movements in bronze corydoras (Corydoras aeneus). Can J Zool 88, 378–389 (2010).

24. Gibb, A. C., Swanson, B. O., Wesp, H., Landels, C. & Liu, C. Development of the Escape Response in Teleost Fishes: Do Ontogenetic Changes Enable Improved Performance? Physiological and Biochemical Zoology 79, 7–19 (2006).

25. White, R. M., et al. Transparent Adult Zebrafish as a Tool for In Vivo Transplantation Analysis. Cell Stem Cell 2, 183–189 (2008).

26. Favre-Bulle, I. A., Vanwalleghem, G., Taylor, M. A., Rubinsztein-Dunlop, H. & Scott, E. K. Cellular-Resolution Imaging of Vestibular Processing across the Larval Zebrafish Brain. Current Biology 28, 3711–3722.e3 (2018).

27. Bianco, I. H. et al. The tangential nucleus controls a gravito-inertial vestibulo-ocular reflex. Current Biology 22, 1285–1295 (2012).

28. Straka, H. & Baker, R. Vestibular blueprint in early vertebrates. Front Neural Circuits 7, (2013).

29. Suwa, H., Gilland, E. & Baker, R. Otolith Ocular Reflex Function of the Tangential Nucleus in Teleost Fish. Ann N Y Acad Sci 871, 1–14 (1999).

30. K. Lee, R. K. & Eaton, R. C. Identifiable reticulospinal neurons of the adult zebrafish, Brachydanio rerio. Journal of Comparative Neurology 304, 34–52 (1991).

31. Stickland, N. C. Growth and development of muscle fibres in the rainbow trout (Salmo gairdneri). J Anat 137 (Pt 2), 323–33 (1983).

32. te Kronnié, G. Axial muscle development in fish. Basic Applied Myology 10, (2000).

33. Jayne, B. C. & Lauder, G. V. How swimming fish use slow and fast muscle fibers: implications for models of vertebrate muscle recruitment. Journal of Comparative Physiology A 175, (1994).

34. Nowroozi, B. N. & Brainerd, E. L. Regional variation in the mechanical properties of the vertebral column during lateral bending in Morone saxatilis. J R Soc Interface 9, 2667–2679 (2012).

35. Tsukamoto, K. Swimming motion of fishes: Their adaptation to water. Hikaku seiri seikagaku(Comparative Physiology and Biochemistry) 10, 249–262 (1993).

36. Hang, H., Heydari, S., Costello, J. H. & Kanso, E. Active tail flexion in concert with passive hydrodynamic forces improves swimming speed and efficiency. J Fluid Mech 932, (2022).

37. Martel, G. & Dill, L. M. Influence of Movement by Coho Salmon (*Oncorhynchus kisutch)* Parr on Their Detection by Common Mergansers (*Mergus merganser)*. Ethology 99, 139–149 (1995).

38. Tang, Z.-H., Huang, Q., Wu, H., Kuang, L. & Fu, S.-J. The behavioral response of prey fish to predators: the role of predator size. PeerJ 5, e3222 (2017).

39. Gerry, S. P. & Ellerby, D. J. Resolving shifting patterns of muscle energy use in swimming fish. PLoS One 9, (2014).

40. McCune, A. R. & Carlson, R. L. Twenty ways to lose your bladder: common natural mutants in zebrafish and widespread convergence of swim bladder loss among teleost fishes. Evol Dev 6, 246–259 (2004).

41. Maina, J. N. & Maloiy, G. M. O. The morphometry of the lung of the African lungfish (*Protopterus aethiopicus*): its structural-functional correlations. Proc R Soc Lond B Biol Sci 224, 399–420 (1985).

42. Baldridge, H. D. Sinking Factors and Average Densities of Florida Sharks as Functions of Liver Buoyancy. Copeia 1970, 744 (1970).

43. Drucker, E. G. & Lauder, G. V. Function of pectoral fins in rainbow trout: behavioral repertoire and hydrodynamic forces. Journal of Experimental Biology 206, 813–826 (2003).

44. Hove, J. R., O’Bryan, L. M., Gordon, M. S., Webb, P. W. & Weihs, D. Boxfishes (Teleostei: Ostraciidae) as a model system for fishes swimming with many fins: kinematics. Journal of Experimental Biology 204, 1459–1471 (2001).

45. Hale, M. E. Developmental Change in the Function of Movement Systems: Transition of the Pectoral Fins between Respiratory and Locomotor Roles in Zebrafish. Integr Comp Biol 54, 238–249 (2014).

46. Westerfield M. The zebrafish book, 5th Edition; A guide for the laboratory use of zebrafish (Danio rerio) University of Oregon Press; Eugene, Oregon: 2007.

47. Harris, C. R. et al. Array programming with NumPy. Nature vol. 585 Preprint at 10.1038/s41586-020-2649-2 (2020).

48. Bradski, G. (2000) The OpenCV Library. Dr. Dobb’s Journal of Software Tools, 120; 122–125.

49. Hunter, J. D. Matplotlib: A 2D graphics environment. Comput Sci Eng 9, (2007).

50. McKinney, W. (2010) Data Structures for Statistical Computing in Python. Proceedings of the 9th Python in Science Conference, Austin, 28 June-3 July 2010, 56-61. 10.25080/Majora-92bf1922-00a.

51. van der Walt S, Schönberger JL, Nunez-Iglesias J, et al (2014) scikit-image: image processing in python. PeerJ 2:e453. 10.7717/peerj.453.

52. Virtanen, P. et al. SciPy 1.0: fundamental algorithms for scientific computing in Python. Nat Methods 17, 261–272 (2020).

