## Supplementary figures and images for "Biomechanics of static roll posture control by body flexion in adult zebrafish"

### Fugure S1

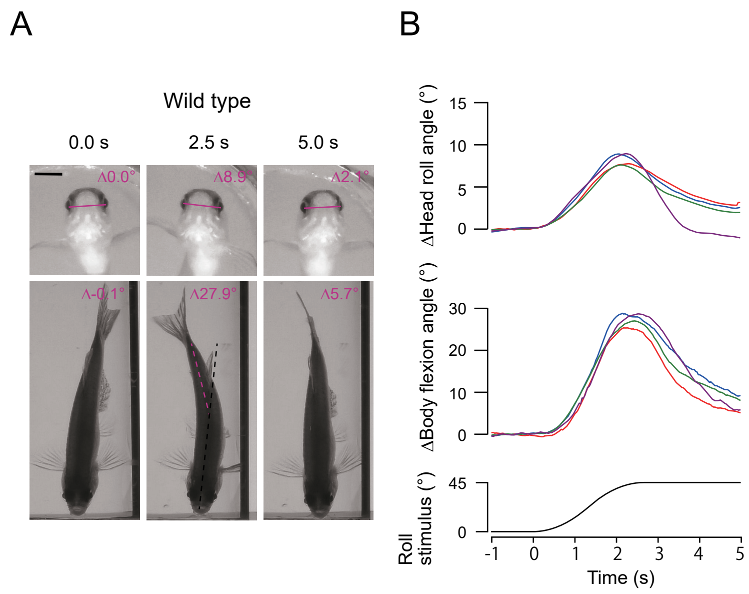

### Fugure S2

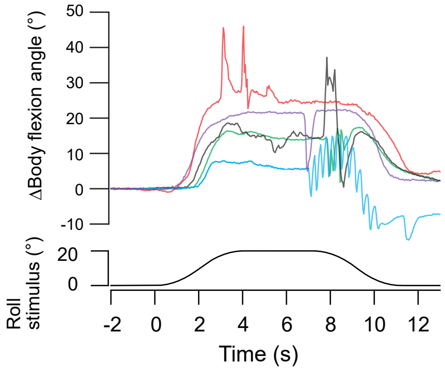

### Fugure S3

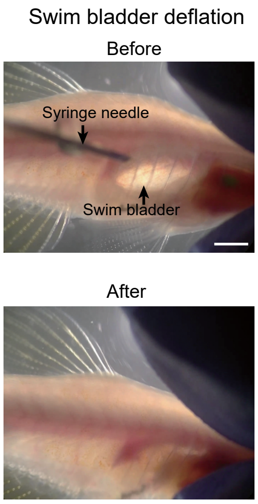

### Fugure S4

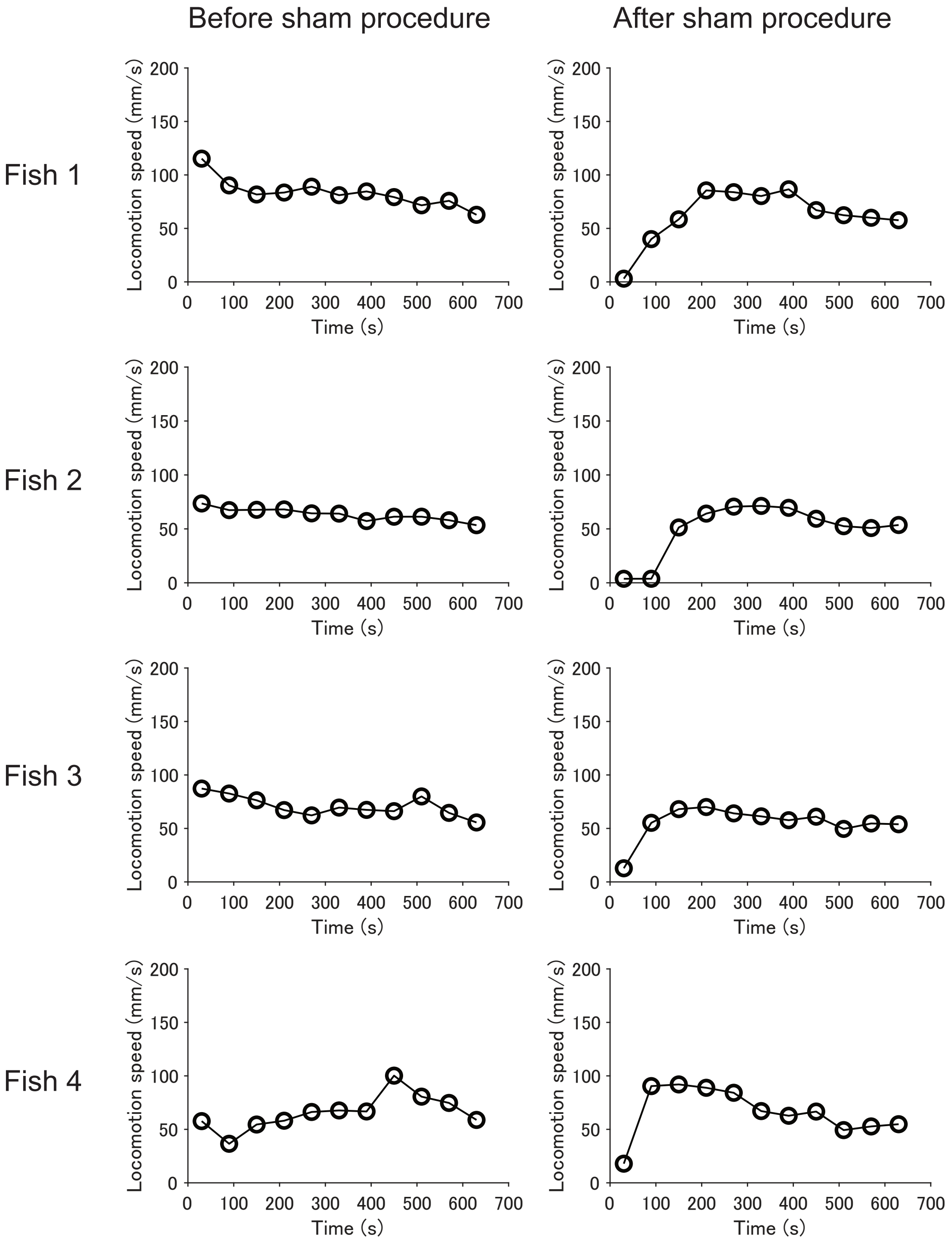
