## Supplementary material for "Biomechanics of static roll posture control by body flexion in adult zebrafish": Table S1

| | | Intact (N=4) | Sham (N=6) | Deflated (N=6) | Intact vs Sham | Sham vs Deflated |
|---|---|---|---|---|---|---|
| Head roll | Onset time (s) | 0.229 ± 0.054 | 0.065 ± 0.052 | 0.176 ± 0.170 | U = 0, *$p$ = 0.014 | U = 8.5, $p$ = 0.145 |
| | Peak time (s) | 2.218 ± 0.186 | 2.024 ± 0.071 | 2.576 ± 0.152 | U = 5, $p$ = 0.171 | U = 0, *$p$ = 0.002 |
| | Maximum angle (°) | 13.147 ± 2.633 | 12.935 ± 2.854 | 19.300 ± 4.990 | U = 11, $p$ = 0.914 | U = 4, *$p$ = 0.026 |
| | Minimum angular velocity (°/s) | -7.466 ± 1.426 | -6.947 ± 1.099 | -4.887 ± 1.009 | U = 15, $p$ = 0.610 | U = 3, *$p$ = 0.015 |
| Body flexion | Onset time (s) | 0.338 ± 0.166 | 0.238 ± 0.058 | 0.454 ± 0.208 | U = 10, $p$ = 0.748 | U = 7, $p$ = 0.093 |
| | Peak time (s) | 2.282 ± 0.105 | 2.087 ± 0.124 | 3.304 ± 0.628 | U = 3, $p$ = 0.067 | U = 1, *$p$ = 0.004 |
| | Maximum angle (°) | 23.239 ± 3.671 | 25.914 ± 3.771 | 26.190 ± 5.276 | U = 15, $p$ = 0.610 | U = 19, $p$ = 0.937 |
| | Minimum angular velocity (°/s) | -9.124 ± 2.218 | -10.230 ± 1.547 | -3.108 ± 1.793 | U = 7, $p$ = 0.352 | U = 0, *$p$ = 0.002 |
